# Circadian entrainment and gating in a natural plant population

**DOI:** 10.64898/2026.01.23.701304

**Authors:** Pirita Paajanen, Tomoaki Muranaka, Luíza Lane de Barros Dantas, Paige E. Panter, Genki Yumoto, Borja Ferrero-Bordera, Mie N. Honjo, Hiroshi Kudoh, Antony N. Dodd

**Author notes:** These authors contributed equally to this work (Pirita Paajanen, Tomoaki Muranaka, Luíza Lane de Barros Dantas, Paige E. Panter).

## Abstract

Circadian clocks coordinate physiology, development and behaviour with cycles of day and night, which contributes to organismal fitness. For this to occur, the circadian clock must be aligned with the natural daily fluctuations in the environmental conditions. Here, we investigated under natural conditions the role of environmental cues in adjusting the circadian clock, and how this affects clock outputs. By combining novel field-based experimentation in a natural plant population with genome-wide analysis and machine-learning strategies for data interpretation, we inferred that temperature cues change circadian timing under natural conditions. We inferred that low temperature cues can shift circadian phase under naturally fluctuating conditions, and that there are seasonal changes in the effects of temperature upon circadian entrainment. We also identified extensive circadian modulation of temperature responses under field conditions. Therefore, plasticity of circadian timing under natural conditions allows flexible responses of both the clock and its outputs to environmental stimuli.

## Introduction

Life on Earth inhabits environments that are generally characterised by daily (24 h) and seasonal cycles. These regular environmental fluctuations have selected for the evolution of circadian clocks. Circadian clocks are cellular mechanisms that produce a biological estimate of the time of day, which organises biological systems over time to confer a selective advantage^1–4^. Circadian clocks generally comprise a biological oscillator formed from molecular feedback loops, which is adjusted by environmental cues to align the timing of the oscillator with 24 h changes in environmental conditions^5^. This adjustment by the environment is known as entrainment, and allows the oscillator to provide an appropriate cellular estimate of the time of day. In turn, output pathways communicate information about the time of day to multiple aspects of physiology, metabolism and behaviour, through the regulation of gene expression and protein activity^5^.

In plants, circadian regulation directly impacts growth rates, responses to environmental stress, the photoperiodic control of flowering, and reproductive fitness^2,6–8^. In natural accessions of the model plant *Arabidopsis thaliana*, there has been selection for latitudinal clines in circadian period length that are thought to be adaptive^9,10^. Furthermore, genetic variation in circadian clock components is associated with crop domestication, and the clock influences traits including flowering time, grain nutrition, and stress tolerance in a variety of crops^11–15^. Because appropriate alignment between the circadian clock and the environmental conditions is necessary for the multitude of benefits derived from clock control^2^, the process of entrainment is of utmost importance for the advantage conferred to plants by circadian regulation. Furthermore, the plant circadian clock engages in extensive modulation of responses to environmental signals^6,16^, which is proposed to ensure that environmental responses are appropriate for the time of day.

Most studies of circadian programs in plants have been conducted under controlled conditions, leading to a gap in understanding of roles for circadian regulation in plants under naturally fluctuating conditions. How do mechanisms and processes identified from model plants under controlled conditions manifest in the field? This is of considerable importance for placing plant circadian clocks into agricultural, ecological and evolutionary contexts, for understanding their adaptive functions, and can reveal novel features of- and roles for- the circadian clock. For example, the effect of far red at the end of the day upon photoperiodic control was identified through field experimentation^17^, as was the effect of crop microenvironments upon circadian clock phase^18^ and the alterations in circadian amplitude caused by seasonal temperature changes outdoors^19,20^.

Here, we investigated the extent to which transient environmental temperature cues can adjust circadian timing in a natural plant population, and how daily (24 h) cycles modulate transcriptional responses to low temperature cues. In naturally fluctuating conditions, daily rhythms of expression of circadian clock genes are remarkably stable across environmental conditions and seasons^19,21^, yet the clock remains responsive to the environment with light conditions and low temperatures during winter and changing the phase and amplitude of circadian rhythms, respectively^18,19^. Furthermore, mathematical modelling of gene expression data from field conditions in rice and *Arabidopsis halleri* predicts that there are 24 h cycles of the sensitivity of some transcripts to the ambient temperature^21,22^. By employing systematic temperature manipulations to plants growing in their natural habitat, we infer that environmental cues can adjust circadian timing under natural conditions.

## Results

Under naturally fluctuating conditions, we investigated the changes in daily timing that are caused by temperature cues, and the daily modulation of transcriptomic responses to transient temperature stimuli. To achieve this, we used a natural population of *Arabidopsis halleri* subsp. *gemmifera* as a model (Fig. 1a, b). This close relative of the circadian clock model *Arabidopsis thaliana* is an evergreen perennial and forms dense clonal patches (Fig. 1a)^23^. We exploited the perennial life history of *A. halleri* to conduct identical experiments during four seasons of the year (Fig. 1b), which exposed the plants to different light and temperature conditions (Fig. 1c, d), to investigate the environmental regulation of the clock and its outputs under natural conditions.

**Figure 1.**
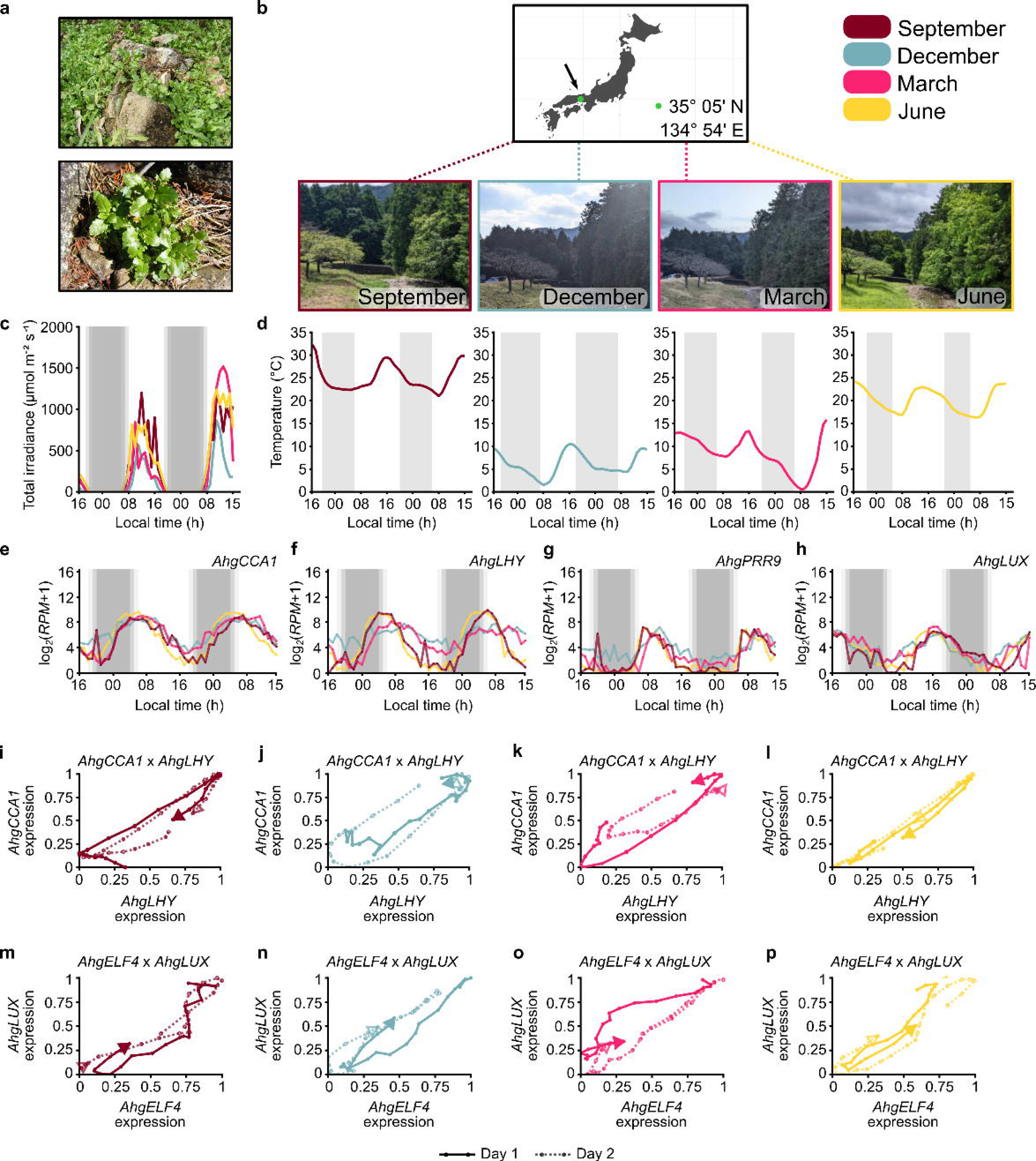
Environmental modulation of circadian oscillator dynamics in a natural plant population. (a) *A. halleri* growing at field site, showing a patch of adjacent plants, and an individual plant. (b) Location of field site and its appearance during four seasons of the year. Diel fluctuation in (c) photon irradiance and (d) ambient temperature at four cardinal points of the year (close to the winter and summer solstice, and the spring and autumn equinox). (e-h) Examples of diel oscillations of circadian oscillator transcript accumulation during four sampling seasons. Grey shading on graphs indicates time between sunset and sunrise (seasonal photoperiod differences shown with transparent shading). (i-p) Temporal alignment between (i-l) *AhgCCA1* and *AhgLHY* transcript accumulation, and (m-p) *AhgELF4* and *AhgLUX* transcript accumulation, over 48 hr sampling periods. In i-p, each graph comprises the normalized abundance of two oscillator transcripts that are plotted against each other, with each data point representing the normalized abundance of both transcripts at a given timepoint. The line joins consecutive data points within the time-series. The solid line represents sampling Day 1 and the broken line represents sampling Day 2. A filled arrow is placed at the end of the time-series from Day 1, and an open arrow is at the end of the Day 2 time-series. Transcripts that are perfectly correlated will produce a diagonal line within the parameter space (e.g. panel l), whereas transcripts that are temporally displaced from each other will produce a plot that rotates through the space (e.g. panel m).

### Plasticity of the circadian oscillator under natural conditions

To investigate the adjustment of the circadian clock by environmental cues in a natural plant population, we required high resolution time-series data to measure subtle changes in oscillator dynamics under field conditions. The oscillator dynamics will be determined by the local environmental conditions at the time of experimentation. Over 48 h periods during four seasons of the year, we collected *A. halleri* leaf tissue for RNA sequencing at 1 h intervals. This identified that, of the transcripts encoding circadian clock components, *AhgCCA1*, *AhgPRR9* and *AhgLUX* had relatively stable oscillations across all seasons (Fig. 1e-h). In comparison, *AhgLHY* had slightly reduced amplitude during the two colder sampling seasons (December, March; Fig. 1f), and *AhgPRR7* was arrhythmic during December and the second day of sampling during March, when temperatures decreased to 0 °C (Fig. S1h). The relatively robust oscillations of most clock components across seasonally-differing environmental conditions (Fig. S1) is consistent with evidence that the oscillator is robust to environmental fluctuations under natural conditions^19,21,24^. Of the oscillator transcripts that were rhythmic (JTK_CYCLE *q* < 0.001), all had a period length near to 24 h (Fig. S1). This was the case also across the entire rhythmic transcriptome (Fig. S2). Furthermore, analysis of the entire transcriptome using a sinusoidal period sweep with least-squares fitting identified a proportion of transcripts that also had an ultradian (< 24 h) oscillation (Fig. S2). This was particularly the case during the March sampling season, when the ambient temperature conditions were quite different on each day of sampling.

Despite this robustness, there were differences in the waveforms of circadian oscillator transcript accumulation during different sampling seasons and days (Fig. 1e-h; Fig S1). This motivated us to quantitatively investigate the environmental modulation of oscillator dynamics under natural conditions. To explore this, we plotted against each other two pairs of circadian oscillator transcripts that are generally regulated with specific phase relationships under controlled conditions in *A. thaliana* (*CCA1* and *LHY*, and *PRR7* and *PRR9*^25^). For example, in an experiment with *A. thaliana* under controlled free running conditions, the oscillation of *CCA1* transcripts occurred slightly in advance of *LHY* (Fig. S3a), and the oscillation of *ELF4* transcripts occurred slightly in advance of *LUX* (Fig. S3b)^25^.

Applying this analysis to the data collected from *A. halleri*, under naturally fluctuating conditions, revealed differences in the temporal relationship between each of these transcript pairs during each sampling season (Fig. 1i-p). The relative accumulation of *AhgLHY* and *AhgCCA1* transcripts had different temporal alignment during each sampling season (Fig. 1i-l) such that during September, *AhgCCA1* was expressed somewhat ahead of *AhgLHY*, somewhat behind during December and March, and concurrently during June (Fig. 1i-l). This suggests that there can be delays or advances in the relative timing of different oscillator components, depending on the environmental conditions. This was also the case for a comparison of *AhgELF4* and *AhgLUX*. During the September and December sampling seasons, the oscillation of *AhgLUX* expression was somewhat advanced relative to *AhgELF4* (Fig. 1m, n). In comparison, during the March sampling, *AhgELF4* was advanced relative to *AhgLUX* on Day 1 of sampling, and they were co-regulated on Day 2 (Fig. 1o; the temperature decreased rapidly on Day 2 of sampling). Furthermore, during the June sampling, *AhgELF4* and *AhgLUX* appeared to have similar phasing (Fig. 1p). Taken together, this shows that the temporal relationship between different circadian oscillator components is flexible, and varies according to fluctuations in the environmental conditions. This is true also at the whole transcriptome scale, where fewer transcripts were rhythmic during the cold sampling seasons of December and March (3153 rhythmic transcripts in September, 941 in December, 1771 in March, 5476 in June (JTK_CYCLE *q* < 0.001)) (Dataset S1).

### Inference of phase changes of oscillator transcripts by exogenous temperature cues under natural conditions

These results suggest that naturally occurring temperature fluctuations change the expression of circadian oscillator components in *A. halleri* over the 24 h cycle. This is consistent with the idea that entrainment of the circadian clock occurs through a process of continuous adjustment of the oscillator in response to the environment, to align its timing appropriately with the environmental conditions^26^. We wished to investigate further the entrainment of the circadian oscillator in a natural plant population, by examining the response of *A. halleri* circadian oscillator transcripts to exogenous environmental stimuli.

Under controlled conditions, entrainment can be studied by measuring the downstream change in circadian oscillator phase that occurs in response to a transient cue applied at several different times of day to different samples^27^. Cues that entrain the clock are often known as zeitgebers (“time giver”). For our experiments, we selected a transient low temperature treatment as a zeitgeber, because transient low temperature treatments alter the phase of both the *A. thaliana* and wheat circadian clocks under controlled conditions in a manner that is consistent with entrainment^28,29^.

We designed bespoke portable equipment to deliver short and consistent cold temperature treatments to *A. halleri* leaves under field conditions (Fig. 2a, b; Fig. S4). Using this, we provided 3 h cold temperature treatments (*ca*. 4 °C) to leaves of different *A. halleri* plants at 1 h intervals, over a 48 h period, alongside ambient-temperature controls (Fig. 2c; Fig. S4). Leaf tissue was harvested for RNA sequencing (RNA-seq) at the end of each 3 h cold treatment, to study alterations in circadian oscillator gene expression that were caused by the cold treatments. To evaluate the effectiveness of cold stimulus delivery, we examined the consistency of the low temperature treatments, which we found to be extremely consistent (Fig. S4). Circadian oscillator transcripts of *A. halleri* underwent changes in expression in response to the cold stimuli, with the changes varying in magnitude according to the time of day (Fig. 2d-f). These responses varied according to the identity of the circadian oscillator transcript (Fig. 2d-f). For example, *AhgPRR9* transcripts were upregulated by a transient cold treatment at times when the transcript levels were declining under ambient temperature conditions (Fig. 2e). In comparison, the oscillator transcript *AhgLUX* was induced in response to transient cold at times prior to its increase under ambient temperature conditions (Fig. 2f).

**Figure 2.**
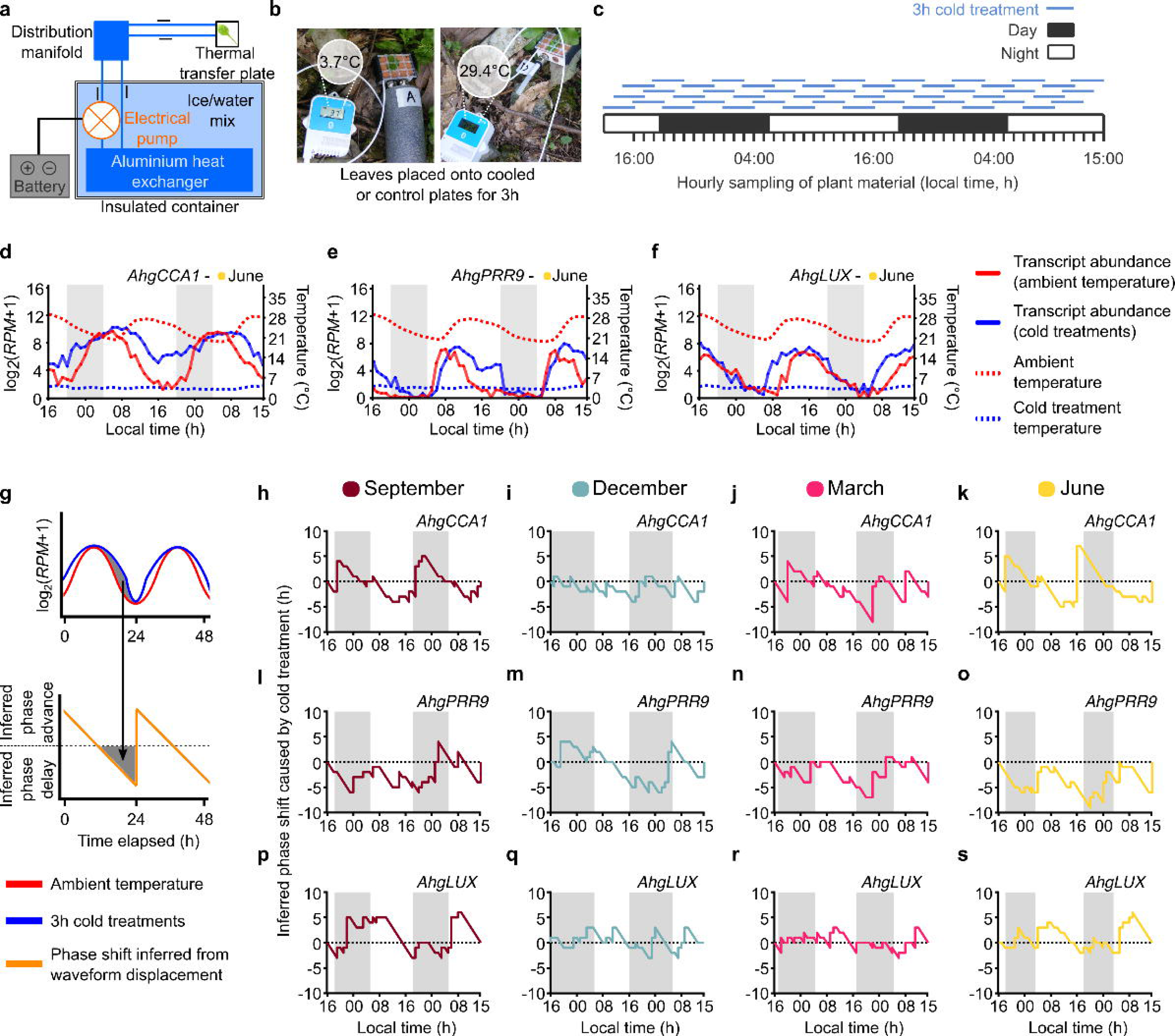
Inference of low temperature entrainment of circadian oscillator components under naturally fluctuating conditions. (a) Apparatus for delivering 3 h cold temperature treatments under field conditions, and (b) its use. The ambient temperature (control) leaf was attached to an identical thermal transfer plate that was not connected to the cooling system. Temperature was measured directly from temperature sensors within the thermal transfer plates. (c) Separate leaves were given 3 h cold treatments staggered by 1 h intervals, over a period of about 48 h, and harvested after each 3 h treatment. Each cold treatment was given to a different plant to avoid resampling effects. (d-f) Transient low temperature treatments alter accumulation of individual circadian oscillator transcripts (exemplified by *AhgCCA1*, *AhgPRR9* and *AhgLUX*). (g) Schematic representation of approach for inference of phase changes at each timepoint that are caused by each transient cold treatment. (h-s) For three circadian oscillator components (*AhgCCA1*, *AhgPRR9*, *AhgLUX*), inferred phase changes local to each timepoint that are caused by short cold treatments at each timepoint over 48 h, across four sampling seasons. Dashed horizontal lines on graphs indicate no inferred phase change, and lines above and below this line represent inferred phase advances and delays, respectively. Grey shading indicates period between sunrise and sunset.

Because transient low temperature treatments altered the abundance of several circadian oscillator transcripts systematically throughout this experiment (e.g. Fig. 2d-f), we reasoned that these small and temporally local alterations might adjust the phase of circadian oscillator components in a manner consistent with entrainment. One classical approach for the investigation of entrainment requires the unperturbed continuous monitoring of the oscillator under free running conditions after each environmental stimulus, to measure the change in phase that is caused by the stimulus^30^. This is not possible under field conditions, where the environment varies continuously. Therefore, we devised a strategy to infer the extent to which each short cold treatment advanced or delayed each circadian oscillator transcript abundance at each timepoint. To achieve this, we inferred the time that it takes for each transcript to reach the same level of expression after the cold treatment, relative to the ambient temperature condition, at each measured point of the daily cycle (in other words, the time taken for the expression level under ambient temperature conditions to catch up with the expression level after the cold treatment, or alternatively for the cold treatment to reach the ambient transcript level). To do this, we used a dynamic time warping algorithm that estimated the minimum distance between the ambient and cold time series^31^. This method accommodates local compressions and stretches in the waveform, and the warping algorithm deforms the cold treated time-series to the ambient time-series. Because the maximum and minimum expression levels for oscillator transcripts were very similar between the cold treated and ambient temperature conditions, the distance between the two time-series was largely in the time scale (*x*) axis, and therefore the difference between the warping provides an estimate of temporal displacement of the expression level for each transcript that was caused by each cold treatment (Fig. 2g). We refer to this as the degree of local entrainment (or phase shift) at each time point, because the entrainment process occurs continuously over the daily cycle^26^.

We applied this analysis to circadian oscillator transcripts that were rhythmic during all sampling seasons (*AhgCCA1*, *AhgPRR9*, *AhgLUX*), because it is not possible to infer phase changes for arrhythmic transcripts. For the oscillator transcripts examined, this approach inferred that cold stimuli applied at differing times in the 24 h cycle can transiently advance or delay the phase of the transcript (Fig. 2h-s). It also inferred that during each 24 h cycle, there was a period when the phase was relatively unresponsive to the stimulus (e.g. Fig. 2h, j, k, n, o, p, s). This dynamic, such as for *AhgCCA1* during the September, March and June sampling seasons (Fig. 2i, j, k), is reminiscent of phase response curves obtained under controlled conditions^32,33^. It is interesting that the timing of the inferred phase shifts varies according to the season of the experiment. During the December and March sampling seasons, *AhgCCA1* (December; Fig. 2i) and *AhgLUX* (December and March; Fig. 2q, r) were inferred to not undergo phase shifts in response to cold temperature stimuli, which might be because the magnitude of temperature difference between ambient conditions and the cold stimulus was insufficient to elicit a response of these circadian oscillator components.

Although cold stimulus-induced phase changes were inferred for *AhgPRR9* during all four sampling seasons (Fig. 2l-o), only phase delays were inferred during the March and June experimentation (Fig. 2n, o). This change alone may not allow correct entrainment of the oscillator, but might be compensated within the oscillator network through competing response(s) of other clock components.

Overall, these inferences suggest that a cold treatment can transiently or continuously shift the phase of these oscillator transcripts under natural conditions, with this response occurring predominantly during warmer seasons. Therefore, the extent to which a temperature stimulus can act as a zeitgeber for the oscillator under natural conditions might depend on the magnitude of temperature alteration. Under controlled conditions, a low temperature pulse (12 °C) can change the phase of several circadian clock reporters in *A. thaliana*^29^, and a pulse of 4 °C can produce changes in the expression of circadian oscillator components in wheat (*T. aestivum*), in a manner reminiscent of entrainment^28^. Furthermore, short periods of increased temperature (27 °C) can also entrain the clock in *A. thaliana*^34^.

### Diel gating of an environmental response under natural conditions

Under controlled conditions, the circadian clock can restrict responses to environmental cues to specific times of day. It can also modulate the magnitude of stimulus-induced responses according to the time of day^6,16,35^. These processes are known as circadian gating, and influence a multitude of environmental responses in plants^6^. It has been inferred using mathematical modelling that there could be gating of environmental responses in plants under naturally fluctuating conditions^19,21^, but this inference has not been tested experimentally. Our experiments provided the opportunity to perform a direct test of the pervasiveness of daily gating of transcriptional responses to an environmental cue in a natural plant population, by evaluating the magnitude of transcriptomic changes following a time-series of cold stimuli across the daily cycle (Fig. 2c).

We adopted an existing conceptual framework that distinguishes two gating types^16^. In the first case (discrete gating), the circadian clock restricts a process to specific time window(s) over the 24 h cycle, so that the response can occur only at certain times (Fig. 3a)^16,35–37^. In the second case (continuous gating), the circadian clock modulates the sensitivity of a process to an environmental cue, such that the magnitude of response is greater at certain times of day than at others (Fig. 3b)^6,16^. Continuous gating is distinct from discrete gating because there is no deactivation of the response at certain times in the 24 h cycle (Fig. 3a, b)^16^. We wished to distinguish between these two phenotypes, and other dynamical aspects of the responses to cold treatments within the time-series. To extend these concepts, we formalized a theoretical categorization of gating types by dividing the waveform into four parts: peak, decreasing, trough and increasing (Fig. 3c). Within this framework, upregulation or downregulation in the response to cold stimuli could theoretically occur within any (or all) of these parts of the waveform, providing 81 theoretical response types, i.e.

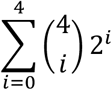

**Figure 3.**
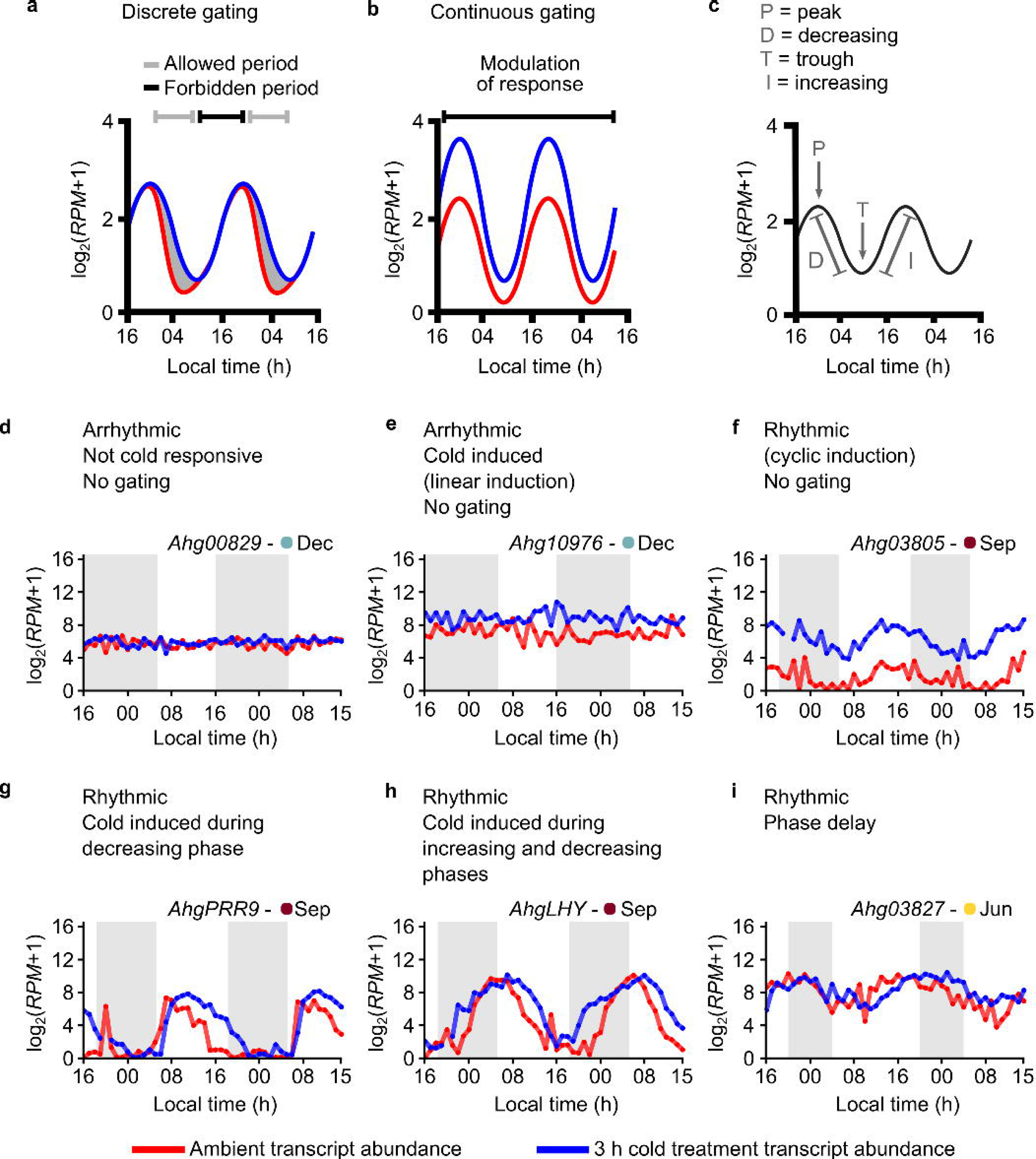
There are diverse temporal patterns of the responses of transcripts to transient low temperature treatments under natural conditions. (a, b) Conceptual representation of discrete gating and continuous gating^16^. (a) During discrete gating, a response to a stimulus is restricted to specific times within the 24 h cycle, and not permitted during other times in the 24 h cycle. (b) During continuous gating, a response to a stimulus is modulated over the 24 h cycle, such that the magnitude of response, itself, has a 24 h cycle. In this mode, there is a rhythm in the sensitivity of the response to the stimulus, but at no times is the response deactivated. (c) Subdivision of the diel oscillation of a hypothetical transcript into segments within which a cold response could occur. (d-i) Responses of six example transcripts to transient cold treatments over a 48 h cycle during the September sampling season. (d-i) Transcripts that do not undergo diel cycles of gating of their cold response. (g-h) Transcripts with several types of gating of their cold response; arrows emphasize time periods and nature of cold responsiveness.

Individual transcripts fit this framework in a variety of ways. Some transcripts did not respond to the cold stimulus (e.g. Fig. 3d), and others responded to the cold stimulus in a similar manner, irrespective of the time of day (e.g. Fig. 3e; upregulation). Other transcripts were rhythmic under ambient temperature conditions and responded to the cold stimulus in an equivalent manner, irrespective of the time of day (e.g. Fig. 3f; upregulated). In all these cases, the transcripts did not have daily rhythms of sensitivity to the cold stimulus and. Therefore, we did not consider such transcripts to have diel cycles of gating of their response to cold.

Within this framework, other transcripts had responses that were consistent with daily gating of cold responsiveness. For example, the circadian oscillator transcript *AhgPRR9* was upregulated by cold during the decreasing phase of its oscillation (Fig. 3g). In comparison, the oscillator transcript *AhgLHY* was upregulated by cold during both increasing and decreasing phases of its oscillation (Fig. 3h). These are both examples of discrete gating (Fig. 3a), because the stimulus-induced response is restricted to specific times of day.

Additional types of response were present, such as upregulation and downregulation at different times in the 24 h cycle, which has the appearance of a phase delay or advance (e.g. Fig. 3i). We consider this to be a form of continuous gating (Fig. 3b) whereby the response can comprise both upregulation and downregulation.

Therefore, in addition to 81 theoretical responses, there were 6 further responses (no expression, no response, phase advance, phase delay, linear induction, linear repression), giving a total of 87 potential responses to the cold stimulus over time. Manual examination of 10% of transcripts found that 23 of these responses occurred experimentally (Fig. S6a-w, with examples for each gating type). Continuous gating manifests as repression and induction during the ascending and descending phases, respectively (Fig. S6h), and nearly all the cold-upregulation gating categories existed in our experiments (e.g. Fig. S6d, f, i-r).

To understand how daily timing influences stimulus-induced responses during different seasons of the year, we wished to identify across the transcriptome the predominant cold-induced gating types during each sampling season. To achieve this, we developed a supervised machine learning model to infer the cold response gating profiles across the transcriptome. This used Tensorflow as a multi-layer-perceptron neural network that includes 3 fully connected hidden layers (Fig. S5a). Some gating classes occurred with a frequency that was too low to train the model, so we combined transcripts that were in different classes, producing a dataset of 9 cold response classes that was sufficiently rich dataset to train the model (Fig. S5b). As its input, the model takes the 96-dimensional data for each transcript (representing the 48 ambient temperature timepoints, and 48 cold treatment timepoints), and has 9 output nodes denoting the gating class inference. The model achieved over 80% accuracy on our test data (Fig. S5c, d), and was applied subsequently the entire dataset. Discrete induction (Fig. 4a) was inferred to be a more common gating type than discrete repression (Fig. 4b), whilst continuous gating was quite rare (Fig. 4c).

**Figure 4.**
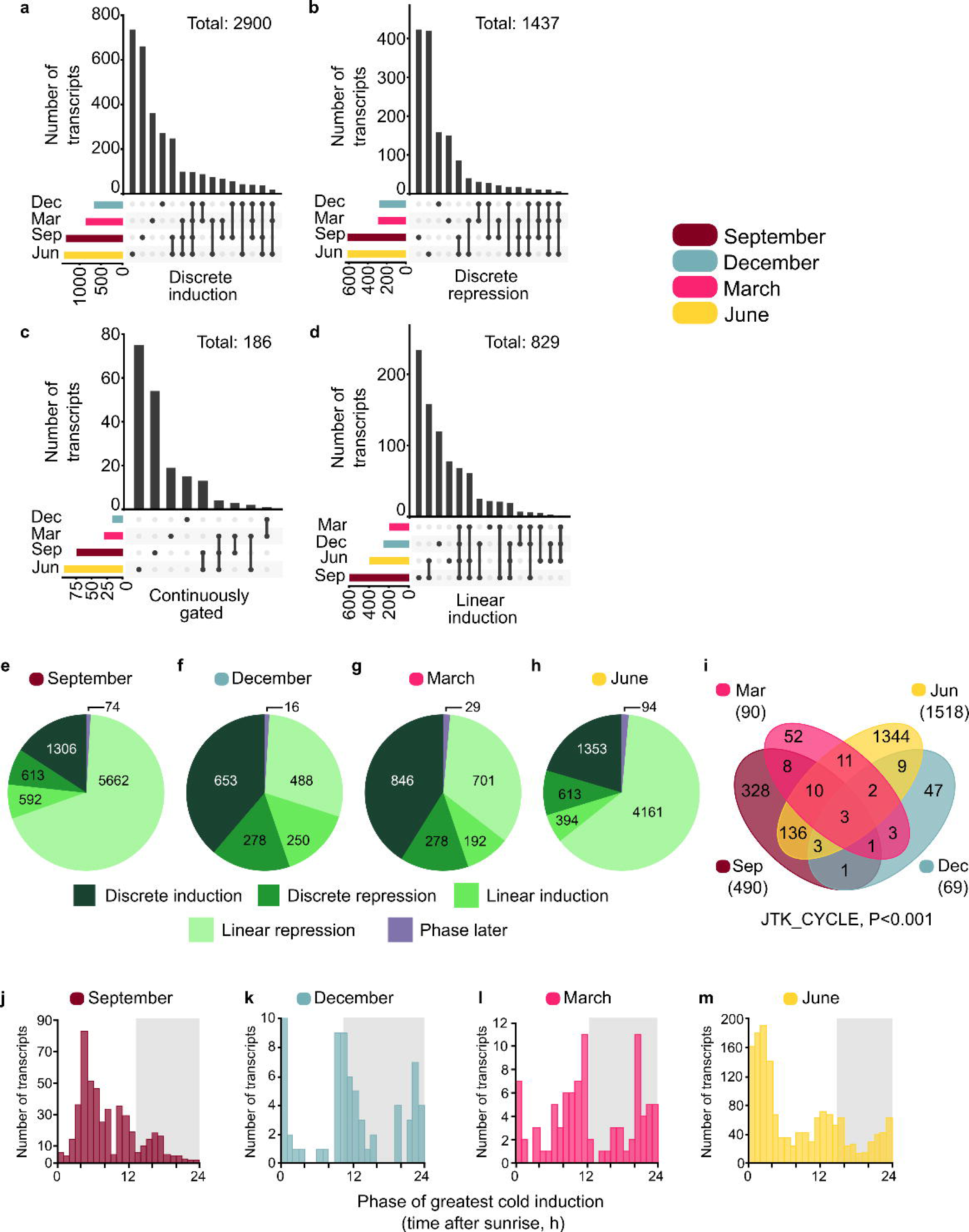
Genome-wide temporal gating of a low temperature response in a natural plant population. (a-d) Numbers of transcripts with each type of major cold temperature response, whereby (a) cold induction and (b) cold repression was restricted to specific times of day, (c) the transcript always responded to cold but with oscillating magnitude of response, and (d) the transcript was always equally cold-responsive, with no effect of time of day on its sensitivity. In (a-d), diagrams under graphs illustrate the transcripts with this temporal response pattern that are unique or common to each sampling season, and coloured horizontal bars indicate the number of transcripts within each class during each sampling season. (e-h) Of transcripts that responded to cold treatments, the proportions of transcripts during each sampling season that had each temporal pattern of cold response (see Fig. 3e). Data are a subset of Fig. S7, with genes removed that were not expressed, not cold responsive, or that had a response that could not be automatically classified. (i) Numbers of transcripts with diel oscillation in cold sensitivity, during all four sampling seasons. (j-m) Phase of greatest cold sensitivity of each transcript undergoing temporal gating of its cold temperature response. Phase was calculated with reference to sunrise time, and grey shading indicates period between sunset and sunrise. The time on x axes is the time relative to sunrise, and grey shading indicates the period between sunset and sunrise.

Remarkably, the number of transcripts that were inferred to have diel gating of their cold response (Fig. 4a-c; 4523 transcripts, 14% of transcriptome) exceeded the number responding equivalently to cold at each time of day (Fig. 4d; 829 transcripts). Up to 37% of the *A. thaliana* transcriptome is rhythmic^25,38–40^. Therefore, the inferred diel cycles of gating of the cold response of 6.1% and 6.3% of all transcripts in September and June, respectively, suggests that 24 h cycles of gating of cold responses is widespread under natural conditions. Of the set of cold responsive transcripts, the cold induction of a greater proportion of transcripts was inferred to be restricted to specific times of day during December and March sampling seasons (i.e. discrete gating). In comparison, during the September and June sampling seasons, the majority of cold induced transcripts were induced by the same magnitude, irrespective of the time of day (Fig. 4e-h; Fig. S7). Our analysis inferred that across the entire transcriptome, the proportion of cold-responsive transcripts was 25% in September, 5% in December, 6% in March and 18% in June, out of 32550 transcripts (Fig. S7). This aligns with the greatest numbers of cold responsive transcripts in *A. thaliana* (around 20% under controlled conditions)^41^.

To determine the time of day of greatest cold sensitivity for each transcript, we calculated for each transcript the difference between its abundance in the cold-treated sample and the ambient temperature control. We then tested whether these time-series of differences had a 24 h rhythms (JTK_CYCLE algorithm)^42^. Transcripts having a statistically significant rhythm (*p* < 0.001) were considered to undergo gating of the response to the low temperature stimulus (Dataset S2). There was variation in the number of transcripts having diel rhythms of cold sensitivity during each sampling season (Fig. 4i), whereby the proportion of transcripts that underwent diel gating of their cold response ranged from 5% in March to 27.7% in June. During the December, March and June sampling, the majority of transcripts had greatest cold sensitivity around either dawn or dusk (Fig. 4j-m), suggesting that greater responses to this stimulus occurred at times when there are predictable transitions in the light conditions. The exception was the September sampling season, where most transcripts reached maximum cold sensitivity close to noon (Fig. 4j). During this sampling season, the night-time temperature always exceeded 20 °C and the daytime temperature started increasing only later in the morning (Fig. 1d). Therefore, during this sampling season, the phase of the greatest number of gated transcripts tended to reflect the ambient temperature.

## Discussion

In this study, we obtained direct evidence that temperature can regulate the timing of expression of circadian clock transcripts in a natural plant population, and inferred that transient low temperature cues can delay or advance the phase of individual circadian clock transcripts. We also obtained the first direct evidence for 24 h cycles of the sensitivity of transcriptome-wide responses to a temperature stimulus in naturally fluctuating conditions. The relatively high sampling frequency that we used also enabled the detection of ultradian oscillations for a small number of transcripts (Fig. S2), with the majority of these being during the March sampling season. One potential explanation for this ultradian oscillation is that the ambient temperature oscillation was very different on the first and second days of sampling during the March season, so perhaps these additional frequencies arise from the combination of clock-controlled oscillations and direct responses to the temperature changes. Our experiments occurred under conditions of entrainment, whereas other studies that identified ultradian harmonics in transcript data from mammals and *Neurospora crassa* were conducted under free running conditions^43,44^.

We focused on inferred phase changes of individual circadian oscillator transcripts in response to transient low temperature cues, rather than overt physiological or behavioural outputs of the plant circadian clock such as photosynthesis or growth rates^45,46^. The network structure of the *A. thaliana* circadian oscillator means that each circadian oscillator transcript responds somewhat differently to specific environmental cues such as light^47^, and some outputs from the circadian oscillator have either greater light- or temperature-responsiveness in their regulation^29^. This creates a set of environmental responses for individual circadian oscillator transcripts that, in combination with the differing phases of expression of each oscillator component, could explain why each oscillator transcript examined responded differently to transient low temperature cues. For example, *AhgCCA1* was inferred to undergo a phase advance in response to a transient low temperature treatment around dusk (Fig. 2h, k), whereas the phase of *AhgPRR9* was inferred to be delayed around this time (Fig. 2l, n, o). We anticipate that these differing responses of individual oscillator transcripts act in combination-across the oscillator network- to establish a particular phase relationship between the oscillator, its outputs, and the naturally fluctuating environment, with this reported to be remarkably stable in field-grown rice^24^.

Our analysis inferred that transient cold stimuli shifted the phase of certain *A. halleri* clock transcripts in only one direction (e.g. *AhgPRR9* during March and June sampling seasons; Fig. 2n, o). In a similar manner, certain entrainment cues also shift the phase of the clock in only one direction in some other organisms^48^. For example, in *Gonyaulax polyhedra*, red light pulses and UV pulses tend to only advance the phase^49,50^, cycloheximide treatments only delay the phase of the *Acetabularia mediterranea* clock^51^, and in one study, light pulses generally only delay the phase of the rhythm of the phototactic response of *Euglena gracilis*^48,52^. Furthermore, radiation-induced DNA damage only advances the mammalian circadian clock^53^ and microbial short chain fatty acids only delay the mammalian peripheral clock at ZT5^54^. This is not dissimilar to synchronization between neurons, whereby neuronal spiking is phase-locked between cells, and within spike time response curves, there are normally only delays or advances in spiking, depending on the measured parameter^55^. Such phase shifts in specific directions for specific circadian oscillator transcripts could represent the tensioning of different clock components against each other to maintain a stable limit cycle for the entire oscillator.

Because our experiments occurred in a naturally fluctuating ecosystem, the data are representative of the conditions at the time of experimentation. Therefore, any experimental repeat conducted during a different season or year will produce different results. The clock transcript *AhgPRR7* provides an excellent example of this, given the proposed role of PRR7 as a signal integration hub for the entrainment of the *A. thaliana* clock^26,56,57^. During the March sampling season, there were contrasting temperature conditions on the two days of experimentation (higher temperatures on Day 1, cold night on Day 2; Fig. 1d), and *AhgPRR7* oscillated on Day 1, but not Day 2. Similarly, in our data and other studies, *AhgPRR7* is arrhythmic during colder winter temperatures (Fig. S1h)^19,20^. This contrasts *AhgCCA1*, which is robustly rhythmic during these sampling periods and in previous reports (Fig. S1a)^21^. Therefore, the differing dynamics of specific oscillator components within the variable natural environment supports the notion that different oscillator components have contrasting rhythmic responses to environmental cues. However, certain conditions may cause bifurcation of the oscillator limit cycle, such as cold temperature conditions leading to arrhythmia of some oscillator components (Fig. S1)^19^. One potential consequence of this may be to restrict the range of stable entrainment of the circadian oscillator to the climate zones to which a plant species is naturally adapted^58,59^.

Our data enable inferences about the temporal relationship between the circadian oscillator and its outputs. Here, we consider the case of *AhgPRR7*, which regulates an array of genes with important roles in development and metabolism (Fig. S8a)^60,61^. *AhgPRR7* is informative because of considerable differences in its oscillatory dynamics during the different sampling seasons (Fig. S1h). We examined the relationship between *AhgPRR7* oscillations and the dynamics of several of its regulatory targets (*LNK1*, *PIF5*, *ABA1* and *AMY3*; identified in *A. thaliana*^61^; Fig. S8b-k). Its gated response to low temperature pulses was evident for its regulatory target *AhgLNK1* (downregulation around dawn, upregulation around dusk; June sampling season; Fig. S8h, i), somewhat evident for *AhgPIF5* and *AhgABA1* (Fig. S8d, e, j, j), and not in *AhgAMY3* (Fig. S8f, g). These regulatory targets all became arrhythmic during the December sampling season, when *AhgPRR7* was arrhythmic (Fig. S8b-k). Furthermore, a simple linear model can explain 86% of the dynamics of clock-associated *AhgLNK1* from the dynamics of *AhgPRR7* (R^2^ = 0.86; p < 0.001), compared to 27% of *AhgPIF5* (R^2^ = 0.27; p < 0.001), 21% of *AhgAMY3* (R^2^ = 0.21; p < 0.001) and 54% of *AhgABA1* (R^2^ = 0.54; p < 0.001; tested with lm function in R). This seems consistent with the notion that greater regulatory distance from the circadian oscillator is associated with the integration of additional layers of information into the control of transcript dynamics, in addition to clock regulation^62^. Interestingly, a recent analysis in *A. halleri* reported that circadian clock oscillations attenuate or lose rhythmicity below approximately 7 °C, and this coincides with the threshold temperature for growth arrest^20^. Our genome-wide identification of diel gating responses provides molecular evidence for how cold responsiveness is temporally-restricted, perhaps to optimize cold acclimation by avoiding unnecessary energy investment in cold responses during times of day when cold exposure of plants is less damaging.

The phasing of clock gene expression in *A. halleri* during the September and June sampling was comparable to the phasing of these transcripts in an experiment conducted with *A. thaliana* in polytunnels^63^. It is unknown whether the regulatory wiring of the oscillator network in *A. halleri* is the same as *A. thaliana*, but the degree of gene homology between the species suggests that the regulation is likely to be similar. Because the *A. thaliana* accession used for the polytunnel experiment is a rapid cycling summer annual accession that flowers rapidly under long photoperiod conditions (Col-0 accession)^63^, it would not have been possible to examine the regulation of its circadian clock under naturally fluctuating conditions throughout all seasons of the year in the same plants. *A. halleri* undergoes vernalization (exposure to prolonged winter cold) that causes epigenetic downregulation of the floral repressor *FLOWERING LOCUS C* (*FLC*), leading to flowering in the spring^64,65^. This is essentially identical to winter annual accessions of *A. thaliana*^66^, with an exception in *A. halleri* being the reactivation of *AhgFLC* within vegetative tissue during late spring and summer, in a manner consistent with its perennial life history^64,67^. In a winter annual (vernalization-requiring) accession of *A. thaliana*, vernalization shortens the free running circadian period^68^, with *PRR7* shifting to a slightly earlier phase under free running conditions after vernalization^69^. Furthermore, *FLC* also participates in temperature compensation of the clock^39^. Because our experiments occurred under conditions of entrainment, such changes in period are unlikely to be detected, although they might contribute to the overall phase relationship between the clock and environment. In *A. thaliana*, there is extensive circadian gating of cold induced transcripts, but this has only been tested under controlled conditions and in rapid cycling accessions^70–72^. A new finding here is that during different seasons of the year, different sets of transcripts undergo diel gating of their response to cold (Fig. 4i). It would be informative in future to determine the extent to which this arises from seasonal changes in chromatin accessibility around either clock genes or genes that are downstream from the clock^65^.

If zeitgebers such as temperature changes originate as environmental features that are stressors or opportunities^73,74^ and circadian regulation provides an adaptation to those challenges, zeitgebers might select for the evolution of circadian clocks. Although competition or cooperation between different zeitgebers is expected to change the dynamics of the circadian oscillator (such as reinforcing, weakening, or shifting the phase of the rhythm), the clock network has evolved to harvest information from certain combinations or sequences of zeitgebers (Fig. S9). One consequence of climate change is that the daily coordination between zeitgebers such as light and temperature will alter (for example, due to relatively warmer nights^75^), and seasonal cues such as temperature and photoperiod will become misaligned^76^. We reason that this will modify the alignment between the clock and environment, with potential impacts upon the adaptive roles of circadian regulation in the natural environment.

## Methods

### Field site and study species

This study was conducted at a field site (Monzen, Taka-cho, Hyogo Prefecture, Japan 35° 05’ N, 134° 54’ E, alt. 145 m^23,77^) that has been used for previous plant molecular ecology studies (Fig. 1b). The study species was *Arabidopsis halleri* subsp. *gemmifera* (Matsum.) O’Kane & Al-Shehbaz (Brassicaceae) (Fig. 1a), which is an evergreen perennial that spreads through both seed production and clonal patch formation. *Arabidopsis halleri* is closely related to *A. thaliana*, with high gene sequence homology and genetic synteny^78^.

This makes it relatively straightforward to identify and annotate candidate genes. Irradiance, temperature, humidity and windspeed were measured by an automated meteorological station located at the field site. For irradiance monitoring, the meteorological station used a LI-COR LI190R quantum sensor covered with a LI-COR LI190R-BX hemispherical diffuser^23^. Temperature measurements were also made for the individual leaves that were sampled (Fig. 1d). Sampling occurred 3-5 September 2019 (sunrise 05:35; sunset 18:27; photoperiod 12 h 52 min), 10-12 December 2019 (sunrise 06:58; sunset 16:50; photoperiod 9 h 52 min), 10-12 March 2020 (sunrise 06:19; sunset 18:05; photoperiod 11 h 46 min) and 2-4 June 2020 (sunrise 04:49; sunset 19:11; photoperiod 14 h 22 min). Field site location map (Fig. 1b) was generated with R version 4.3.1 running the maps 3.4.2.1 package.

### Transient cold temperature treatments

We constructed a portable apparatus to deliver low temperature treatments to leaves under field conditions. This comprised a large insulated box that contained an ice/water mix, into which was fitted a custom-made heat exchanger (Fig. 2a). Water was circulated through the heat exchanger by a car battery-powered pump that supplied the chilled water through insulated pipework to small thermal transfer pads onto which individual leaves were appressed (Fig. 2b). Leaves were also appressed to control thermal transfer pads that were not supplied with chilled water (Fig. 2b). Temperature loggers (Omni Instruments TR42 Bluetooth temperature logger) with fine temperature sensing probes were incorporated into the thermal transfer pads to track the treatment and control temperatures during the experiment. The entire apparatus was monitored continuously during experiments to ensure stability of the low temperature treatments. During all seasons of experimentation, the apparatus consistently delivered cold temperature treatments to leaves of about 3-5 °C.

### Tissue sampling

All leaf tissue was sampled from a single very large patch of a great number of adjacent *A. halleri* plants at the field site. Leaves were selected for sampling on the basis that they were not overlapped (shaded) by leaves of the same or adjacent *A. halleri* plants, or by other nearby vegetation. The large number of plants at the sampling location allowed each plant to be sampled only once, to avoid any transcriptomic or oscillator changes caused by resampling of the same plant. We always sampled a fully expanded leaf that was not senescent, not showing signs of insect or other damage, and did not have obvious anthocyanin accumulation that can be diagnostic for abiotic or biotic stress.

For each cold treatment timepoint, an *A. halleri* leaf was appressed against a chilled thermal transfer pad, and secured in place with soft mesh. In parallel, a control leaf from a separate but nearby plant was appressed against a thermal transfer pad that was not chilled. After the 3 h cold treatment, the treatment leaves were detached with ethanol-washed dissecting scissors and placed into an Eppendorf tube containing 500 μl of RNALater RNA preservation fluid (Sigma-Aldrich). Tissue was kept on ice at the field site, transferred to a portable -50 °C freezer at 6 h intervals, and stored at -80 °C after completion of the experiment. Each individual plant was sampled only once. We favoured an experimental design that involved sampling a single leaf at hourly timepoints, from different plants within the same clonal patch, because this provides better phase, period and rhythmicity estimation compared with greater replication at more widely spaced timepoints^79–81^.

### RNA-seq library preparation

Leaf samples were homogenised in lysis/binding buffer using a multi-beads shocker (Yasui Kikai, Osaka, Japan). The mRNA was isolated directly from the homogenate using streptavidin magnetic beads (New England Biolabs, Ipswich, MA, USA, #S1420S) and 5ʹ biotinylated polyT oligonucleotide. RNA libraries were prepared using the Breath Adapter Directional sequencing (BrAD-seq) method for strand-specific 3ʹ digital gene expression quantification^82^. Briefly, the mRNA was heat-fragmented and primed with a 3ʹ adapter-containing oligonucleotide primer targeting the polyA tail of the mRNA. The cDNA was synthesised using RevertAid Reverse Transcriptase (Thermo Fisher Scientific, #EP0441) using a Veriti Dx Thermal Cycler (Thermo Fisher Scientific). The 5ʹ adapter was added by strand-specific breath capture and the second strand was synthesised using DNA Polymerase I (Thermo Fisher Scientific, #EP0041). Final PCR enrichment was performed using oligonucleotides containing the full adapter sequence with a unique index for each sample. The PCR products were purified and size selected using AMpure XP beads (Beckman Coulter, Brea, CA, USA, #A63881). The size distribution and concentration of the library were measured using a Model 2100 Bioanalyzer (Agilent Technology, Palo Alto, CA, USA) and QuantiFluor DNA System (Promega, Madison, WI, USA) with an Infinite 200 PRO microplate photometer (TECAN, Basel, Switzerland), respectively. Products from 48 samples were pooled as a library for Illumina sequencing systems. The two libraries were sequenced using two lanes on a HiSeq 2500 instrument (Illumina). The original BrAD-Seq protocol was modified to use KAPA HiFi HotStart ReadyMix (Kapa Biosystems, Woburn, MA, USA, #KK2062) for the final PCR^83^.

### RNA-seq data analysis

The 50 base single-end reads with index sequences were determined using the HiSeq 2500 (Illumina) with the TruSeq v3 platform. Pre-processing and quality filtering were performed using trimmomatic-v0.36. Reference sequences used were nuclear and chloroplast transcript sequences of *A. halleri*, 8,109 viral sequences (NCBI GenBank), and ERCC spike-in control (Thermo Fisher Scientific). Transcripts of *A. halleri* (32,550 genes) were annotated using BLAST best hit against Araport11^84^. The pre-processed RNA-seq reads were mapped to the references and quantified using RSEM 1.2.31 and Bowtie2 2.3.4.1. The estimated read count of each gene was converted to log_2_ (rpm + 1). Mapping between *A. halleri* gene locus codes and *A. thaliana* gene annotations was conducted according to Dataset S4.

### Rhythmic transcript detection

For determination of rhythmicity and quantification of rhythmicity parameters, we used the package MetaCycle 1.2.0^85^ in R 4.2.1 (2022-06-23)^86^. We analysed rhythmicity using the JTK_CYCLE algorithm in MetaCycle (p-value < 0.001 cutoff for rhythmicity)^42^. This same method was applied to determine rhythmicity and phase of the pointwise differences between the cold treated and ambient samples, again using a p-value < 0.001 to call rhythmicity. The phase estimates were derived from JTK_CYCLE analysis.

### Period length distribution across transcriptomes

Period estimation was performed by fitting sinusoidal models across a dense range of candidate periods (4–48 h). For each candidate period, the sinusoidal coefficients were estimated by linear least-squares regression, and the period corresponding to the lowest residual sum of squares was selected. Only transcripts with a coefficient of determination (R^2^) higher than 0.3 were considered.

### Phase shift inference

Low temperature-induced phase shifts were inferred at each timepoint with an approach that compared the time-series collected under ambient temperature conditions with the time-series of low temperature treatments. The transcript data of both the ambient and cold-treated samples for each gene was smoothed by taking a rolling average of the five neighbouring time points in R using package zoo 1.8.12^87^. The smoothed ambient and cold-treated curves were aligned using dynamic time warping (DTW) algorithms from the R package dtw 1.23.1, using default parameters^31^. DTW is a technique for comparing time-series^88^. It provides a distance (time) measure between the time-series that is insensitive to local compression and stretches of the time-series, with the warping deforming one of the time-series onto the other. We used DTW to warp the time-series of low-temperature treated samples into the ambient time series, and calculated the time-variable (x-axis displacement) of the warping to obtain the temporally local phase response plots (Fig. 2g).

The method works as follows. Let the ambient measurements in season *S* ∈ *{Sep, Dec, Mar, Jun}* be called *A_S_* = (*a*_1_*, a*_2_*, …, a*_48_) be the reference time series, and the cold measurements *C_S_* = (*c*_1_*, c*_2_*, …, c*_48_) be the test or query time series. Each measurement *a_i_* consists of the time in hours *i* and the corresponding transcript expression level *e_i_*, and is indexed in *i* = 1*, …,* 48 and similarly for the cold measurement *c_j_* are indexed by *j* = 1*, …,* 48. There are 48 measurements because each time series (ambient temperature and cold temperature) had 48 timepoints.

We now create a 48 *×* 48 matrix with entries:

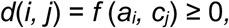

where we can choose *f* to be the Euclidean distance function. Then, we wish to find two warping functions, *ϕ_A_* and *ϕ_C_*, which relate the indices of the two expression levels *A_S_* and *C_S_* to a common time axis *k:*

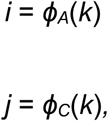

where *k* = 1, 2*, …, T* .

We then wish to find a global pattern dissimilarity measure *D_ϕ_*(*A_S_, C_S_*) where *ϕ* = (*ϕ_A_, ϕ_C_*) are the warping functions. The aim is to find the warping functions that minimize the dissimilarity measure, which is accomplished with the DTW R package^31^.

After the warping functions are found that minimise the *D_ϕ_*(*A_S_, C_S_*), we use the warping matrix and take *ϕ_A_*(*k*) as the new index for time, as a replacement of *i*, and calculate how much the warping has displaced *j*, which is the query-in our case the cold treated sample-to can calculate the new *j* = *ϕ_C_*(*k*) *− ϕ_A_*(*k*), for *k* = 1*, …,* 48. The local phase response curve is then mapping these new *i* and *j*, both now measured in time, i.e., (*ϕ_A_*(*k*)*, ϕ_C_*(*k*) *− ϕ_A_*(*k*)) for *k* = 1*, …,* 48. This gives the time axis measurement, because the maximum and minimum expression levels stay very similar between the cold treated and ambient samples.

### Machine learning model design, features and validation

We used ggplot2 to create plots of 32550 *A. halleri* transcripts, plotting both the ambient and cold-treated timeseries on the same plot for all the four seasons. From this set, we randomly selected 5081 genes for manual classification. Because this random set did not contain enough cyclic genes, we added 1544 known cold induced genes and 1552 rhythmic genes to the classification. This led to 8277 transcripts, which is 6.4% of the transcriptome when multiplied over all four seasons of experimentation. Each plot was inspected and assigned to a gating class using PyQt Image Annotation Tool (https://github.com/robertbrada/PyQt-image-annotation-tool). The final set of genes that were classified and used for model training are in Dataset S3. Machine learning was conducted using TensorFlow 2.15.0 within the package Keras 2.15.0 in Python 3.9.6. The computational details of this are in the GitHub repository for this study. The output of the machine learning was filtered so that the class assigned to each transcript with the greatest probability was assigned as its class; however, if this maximum probability was less than 0.7, then the transcript was called unclassified. This filtering reduced the noise, because some transcripts that did not fall into clear timeseries.

We found that the initial set of 23 gating classes was too sparse for model training, so we reduced the number of gating classes to 3 (discrete induction, discrete repression and continuous gating), with 4 other cold responses (linear induction, linear repression, cyclic induction, cyclic repression), and classes for no expression and no cold response (Fig. 3e). Continuous gating was represented as a “phase later” category (Fig. 3e). Although cyclic induction (Fig. S6f) and cyclic repression (Fig. S6g) occurred for only a small number of transcripts (in our training data, 7 and 12, respectively), we included these for biological completeness. A large number of transcripts underwent “linear repression” (Fig. 3e) (5662 and 4161 during September and June sampling seasons, respectively; Fig. S7). The magnitude of repression was so small that these differences are unlikely to be identified using conventional fold-change differential gene expression analysis, whereas these responses were detected using this machine learning model. However, because that this change often occurred consistently across 48 time points, the interpretation appears well-evidenced, demonstrating that machine learning analysis of time-series data can identify features that would otherwise be missed.

## Data availability

The RNASeq data for this study are freely available in the European Nucleotide Archive (ENA; https://www.ebi.ac.uk/ena) and will be available upon publication with the project ID PRJEB83648. The large datasets within this study, including Datasets S1-S4, are deposited at Zenodo (doi:10.5281/zenodo.14811107).

## Code availability

All code used in this project is available at https://github.com/paajanen/Ahalleri_field_study

## Author contributions

AND and HK conceived the study; TM, PEP, GY, MNH, HK, AND conducted experiments and designed equipment; PP, HK, AND formulated analytical strategies and conceptual development; PP, PEP, TM, BFB, HK, AND analysed and interpreted data; PP built models; PP, LLBD, PEP, TM, GY, MNH, HK, AND designed and wrote the paper.

## Acknowledgements

We thank Richard Morris, Jacqueline Kimmey, Azam Lashkari, Dolf Weijers, Martha Merrow, María Verónica Arana, Carlos Hotta, Marcelo Yanovsky and Stephanie Williams for helpful feedback. Figures 2g, 3ab, S5b, S8a, S9 generated with biorender.com.

## Supplementary figure legends

**Figure S1.** Transcript abundance of circadian oscillator-associated transcripts, across entire timecourses and all four sampling seasons. Grey shading on graphs indicates the time periods between sunset and sunrise. Estimated period lengths are indicated above each graph, with NR indicating datasets that were not called as rhythmic (JTK_CYCLE *q* < 0.001).

**Figure S2.** Distribution of best-fitting periods for all rhythmic transcripts obtained by sinusoidal least-squares period fitting during each sampling season. Data are for samples obtained under ambient temperature conditions. Data are for samples obtained under ambient temperature conditions.

**Figure S3.** Temporal relationship between two transcript pairs under controlled free running conditions. Data were collected under conditions of constant light and temperature, with data deriving from Romanowski et al. 2020^25^ and having a 2 h sampling interval over 48 h^25^. Each graph comprises the normalized abundance of two oscillator transcripts that are plotted against each other, with each data point representing the normalized abundance of both transcripts at a given timepoint. The line joins consecutive data points within the time-series. The solid line represents Day 1 and the broken line represents Day 2. A filled arrow is placed at the end of the time-series from Day 1, and an open arrow is at the end of the Day 2 time-series. Transcripts that are perfectly correlated will produce a diagonal line within the parameter space, whereas transcripts that are temporally displaced from each other will produce a plot that rotates through the space (e.g. panel m).

**Figure S4.** Temperature conditions of individual leaves under field conditions. Comparison of control (ambient) leaf temperature and cold temperature treatment of leaves during each sampling season. Temperature was measured directly from the apparatus used to deliver 3 h cold temperature treatments to individual leaves.

**Figure S5.** Development and structure of machine learning model for interpretation of gating responses. (a) Neural net structure used for classification of diverse temporal cold-temperature dynamics across transcriptome. (b) Summary of major temporal categories of cold-responsiveness present within whole-transcriptome data that were sufficiently abundant to train the machine learning model. (c) Convergence of machine learning model for identification of gating responses to cold. (d) Excellent classification of temporal gating patterns by machine learning model. Coloured scale indicates numbers of transcripts.

**Figure S6.** Diversity of temporal gating of cold temperature responses in *A. halleri* under naturally fluctuating conditions. Each time-series graph represents a distinct category of cold temperature response. The main panel of each is the transcript abundance under ambient temperature conditions and following a 3 h cold temperature treatment. The small (upper right) diagram next to each panel provides a theoretical representation of the class of response. Abbreviations for part of the waveform during which a cold temperature response can occur are P (peak), D (decreasing), T (trough) and I (increasing) waveform sections (illustrated in Fig. 3c). Gene locus IDs shown in italic. Grey shading on graphs represents the time periods between sunset and sunrise.

**Figure S7.** Distribution of temporal gating classes across the transcriptome during each sampling season. Unclassified transcripts are those that could not be assigned to a specific class with at least 70% probability, and represent transcripts such as those responding rapidly to transient environmental fluctuations.

**Figure S8.** Temporal relationship between a clock component and its regulatory targets under naturally fluctuating conditions. This example considers the case of AhgPRR7. (a) Simplified diagram of parts of the circadian oscillator network from *A. thaliana* (left) and some identified regulatory targets of PRR7 in *A. thaliana* (right, purple). (b-j) For two sampling seasons, time-series of (b, c) *AhgPRR7* transcripts and its regulatory targets (d, e) AhgABA1, (f, g) AhgAMY3, (h, i) AhgLNK1, a circadian oscillator associated component, and (j, k) AhgPIF5. See Discussion for explanations.

**Figure S9.** Concept of the gating of circadian clock outputs in the presence of competing zeitgebers. The circadian clock integrates various environmental cues that it receives, using these either for entrainment, to gate environmental signalling, or both. During the morning, Component A receives light and temperature information. For this clock component, light information is the prevailing zeitgeber, while clock outputs have temperature responses that are gated by the clock. The entrainment of Component A affects the circadian phase of Component B, which is entrained predominantly by temperature rather than light conditions, in turn affecting Component A until this is again regulated by light. Both Component A and Component B can be involved in circadian gating of the environmental responses of other transcripts, where the roles of Component A and Component B may differ in producing distinct gating classes. The hypothetical gating class of an output gene from Component A follows its response of A to the temperature cue (producing no clock phase shift), whereas the gating class of an output gene gated by Component B may receive the same phase shift as Component B does, due to its entrainment to the temperature cue. Sun and wave icons indicate light and temperature cues, respectively. Grey arrow thickness indicates the effectiveness of each cue in regulating each of Component A and B. Time-series diagrams at bottom indicate distinct gating classes.

**Dataset S1.** Transcripts with 24 h cycles in abundance under ambient temperature conditions, in a natural population of *A. halleri*. These time series were analysed using MetaCycle (JTK_CYCLE algorithm^85^), to identify transcripts having a significant 24 h oscillation in abundance, and establish their peak phase of expression.

**Dataset S2.** Transcripts called as having a 24 h cycle in their response to a low temperature treatment. The difference in transcript abundance between ambient temperature conditions and following a 3 h cold treatment was calculated for each transcript, at each timepoint. These time-series of differences were analysed using MetaCycle (JTK_CYCLE ^85^) to evaluate whether there was a 24 h rhythm in the magnitude of the response of the transcript abundance to cold, for each transcript. Analysis was conducted separately for the data collected during each sampling season.

**Dataset S3.** Training dataset for machine learning model that was developed to classify 24 h cycles of gating of cold temperature responses within time-series transcriptome data from *A. halleri*.

**Dataset S4.** Mapping of *A. halleri* gene locus coding to *A. thaliana* locus codes and gene names.

