## Supplementary Figures 1-9 for "Circadian entrainment and gating in a natural plant population"

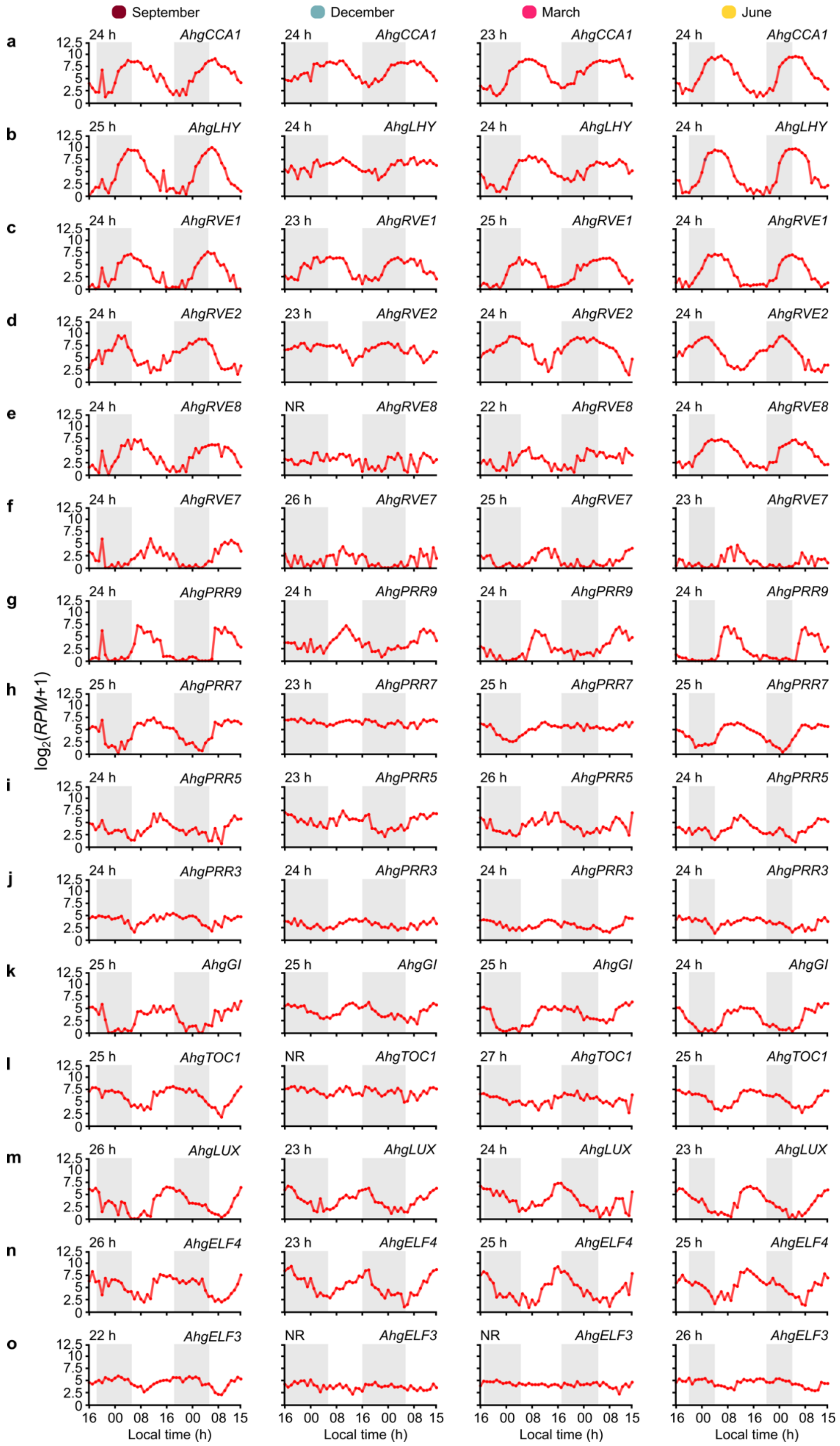


**Figure S1.** Transcript abundance of circadian oscillator-associated transcripts, across entire timecourses and all four sampling seasons. Grey shading on graphs indicates the time periods between sunset and sunrise. Estimated period lengths are indicated above each graph, with NR indicating datasets that were not called as rhythmic (JTK_CYCLE $q < 0.001$).

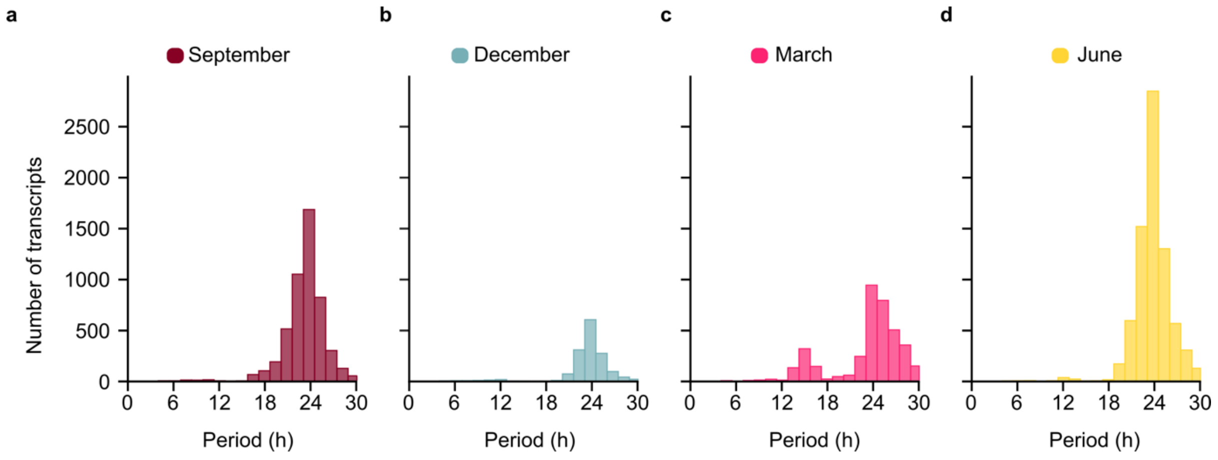


**Figure S2.** Distribution of best-fitting periods for all rhythmic transcripts obtained by sinusoidal least-squares period fitting during each sampling season. Data are for samples obtained under ambient temperature conditions. Data are for samples obtained under ambient temperature conditions.

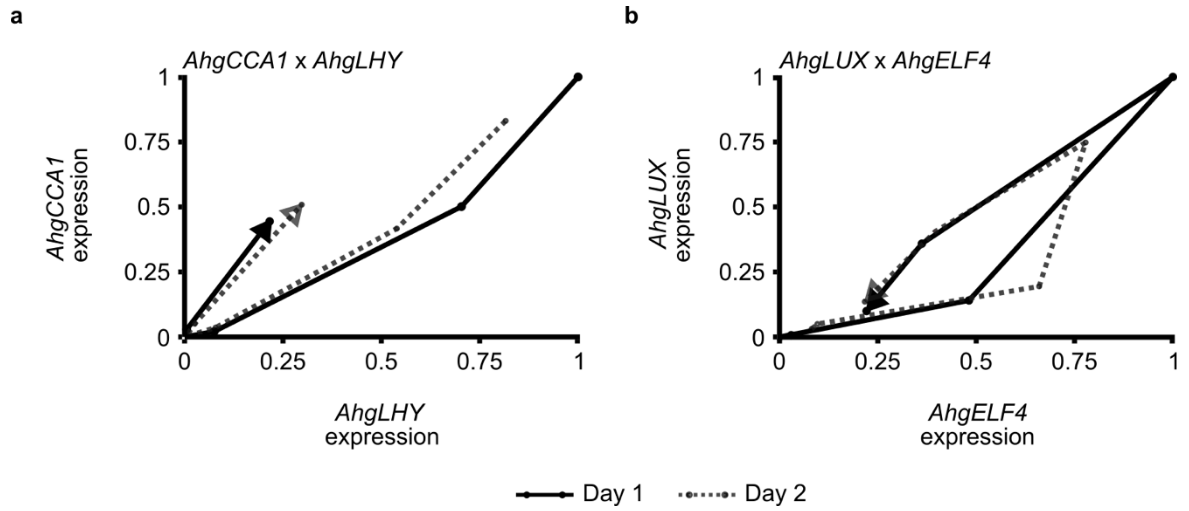


**Figure S3.** Temporal relationship between two transcript pairs under controlled free running conditions. Data were collected under conditions of constant light and temperature, with data deriving from Romanowski et al. 2020[1] and having a 2 h sampling interval over 48 h. Each graph comprises the normalized abundance of two oscillator transcripts that are plotted against each other, with each data point representing the normalized abundance of both transcripts at a given timepoint. The line joins consecutive data points within the time-series. The solid line represents Day 1 and the broken line represents Day 2. A filled arrow is placed at the end of the time-series from Day 1, and an open arrow is at the end of the Day 2 time-series. Transcripts that are perfectly correlated will produce a diagonal line within the parameter space, whereas transcripts that are temporally displaced from each other will produce a plot that rotates through the space (e.g. panel m).

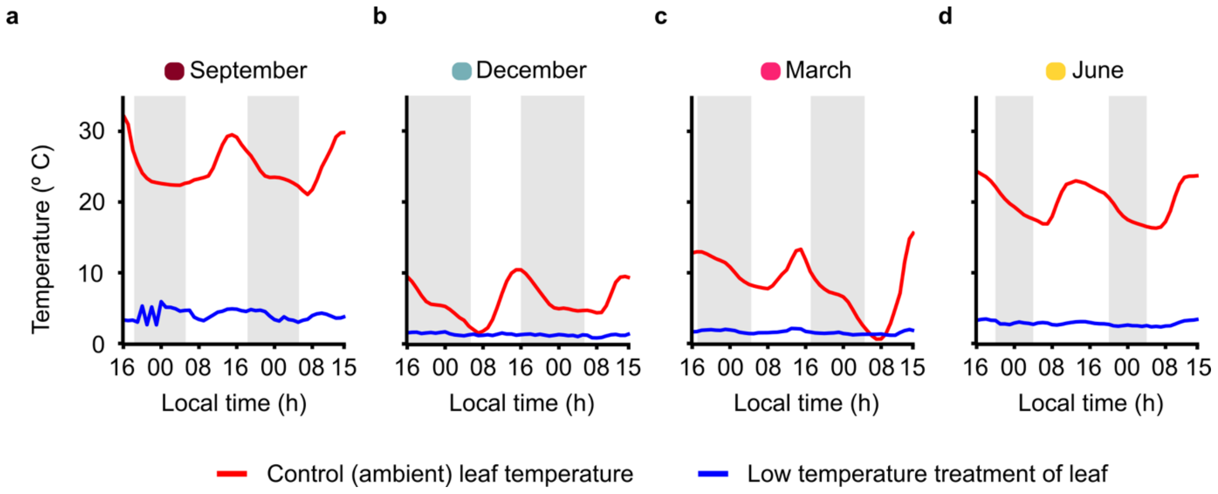


**Figure S4.** Temperature conditions of individual leaves under field conditions. Comparison of control (ambient) leaf temperature and cold temperature treatment of leaves during each sampling season. Temperature was measured directly from the apparatus used to deliver 3 h cold temperature treatments to individual leaves.

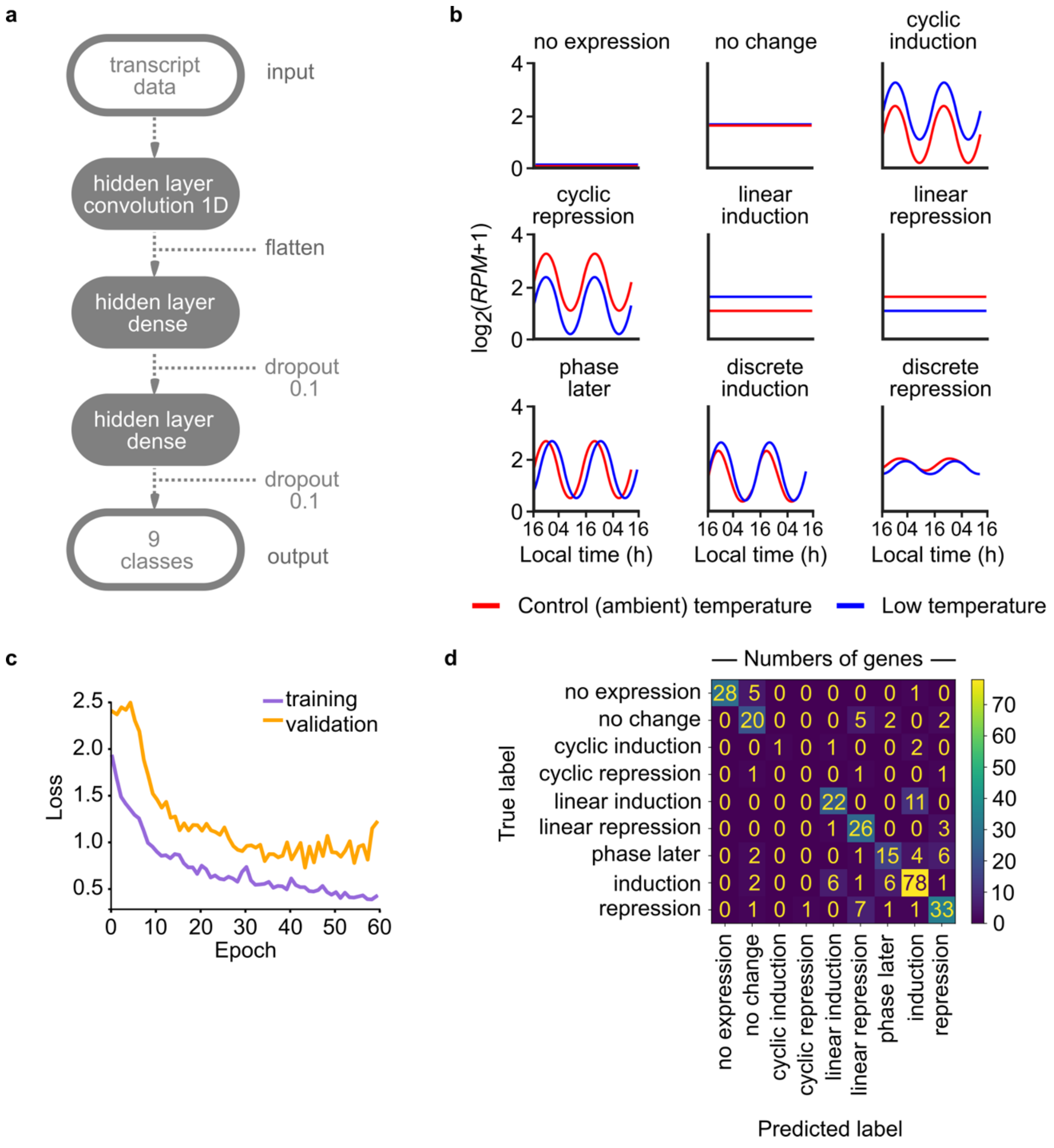


**Figure S5.** Development and structure of machine learning model for interpretation of gating responses**.** (a) Neural net structure used for classification of diverse temporal cold-temperature dynamics across transcriptome. (b) Summary of major temporal categories of cold-responsiveness present within whole-transcriptome data that were sufficiently abundant to train the machine learning model. (c) Convergence of machine learning model for identification of gating responses to cold. (d) Excellent classification of temporal gating patterns by machine learning model. Coloured scale indicates numbers of transcripts.

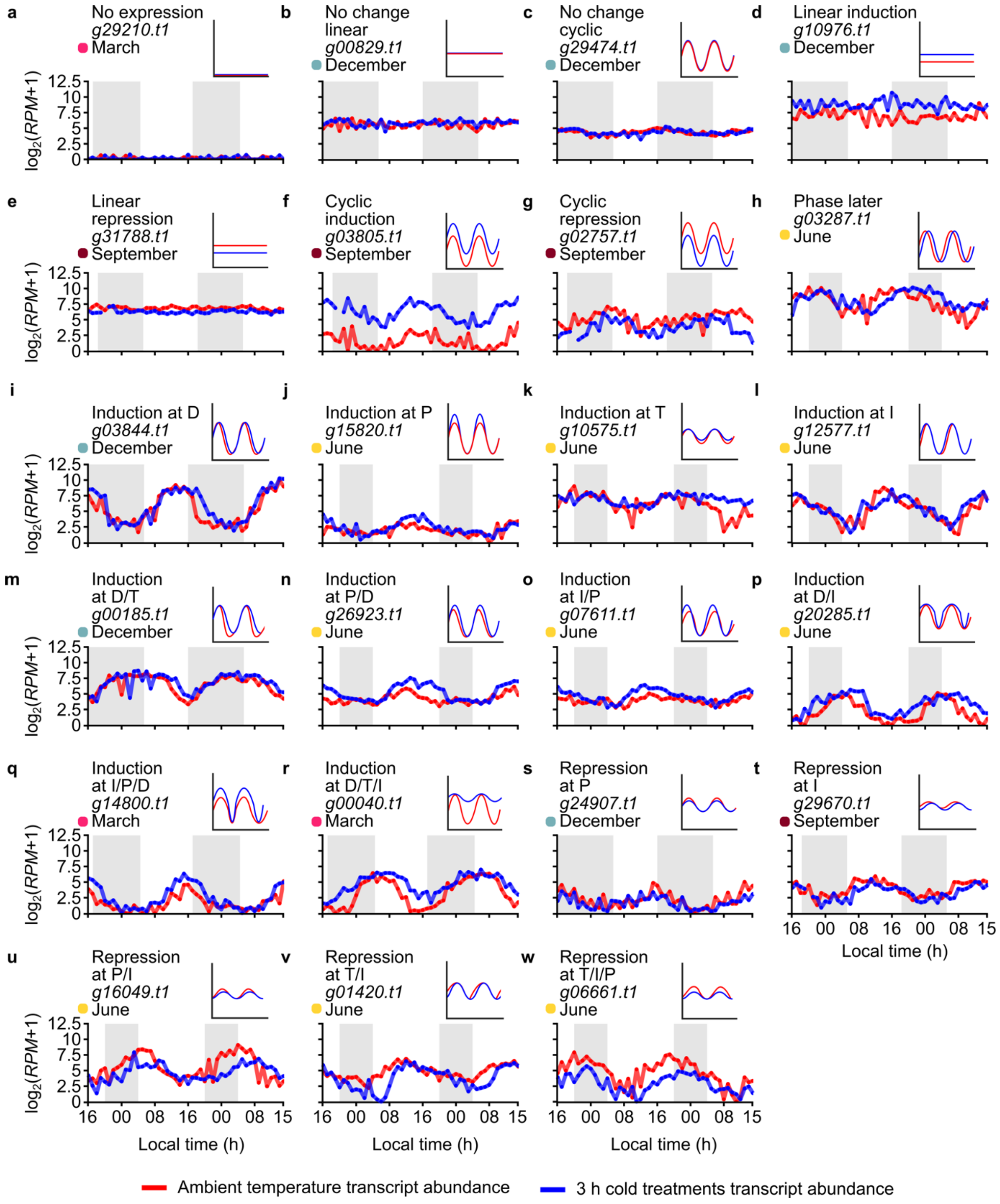


**Figure S6.** Diversity of temporal gating of cold temperature responses in *A. halleri* under naturally fluctuating conditions. Each time-series graph represents a distinct category of cold temperature response. The main panel of each is the transcript abundance under ambient temperature conditions and following a 3 h cold temperature treatment. The small (upper right) diagram next to each panel provides a theoretical representation of the class of response. Abbreviations for part of the waveform during which a cold temperature response can occur are P (peak), D (decreasing), T (trough) and I (increasing) waveform sections (illustrated in Fig. 3c). Gene locus IDs shown in italic. Grey shading on graphs represents the time periods between sunset and sunrise.

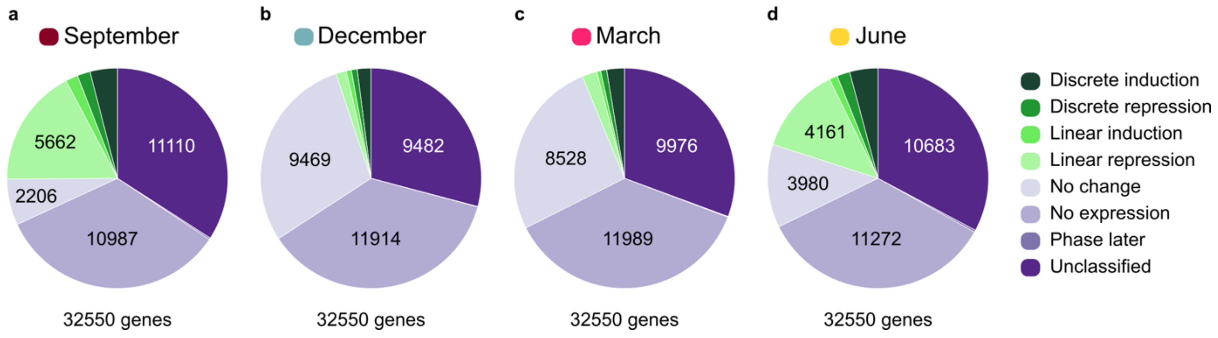


**Figure S7.** Distribution of temporal gating classes across the transcriptome during each sampling season. Unclassified transcripts are those that could not be assigned to a specific class with at least 70% probability, and represent transcripts such as those responding rapidly to transient environmental fluctuations.

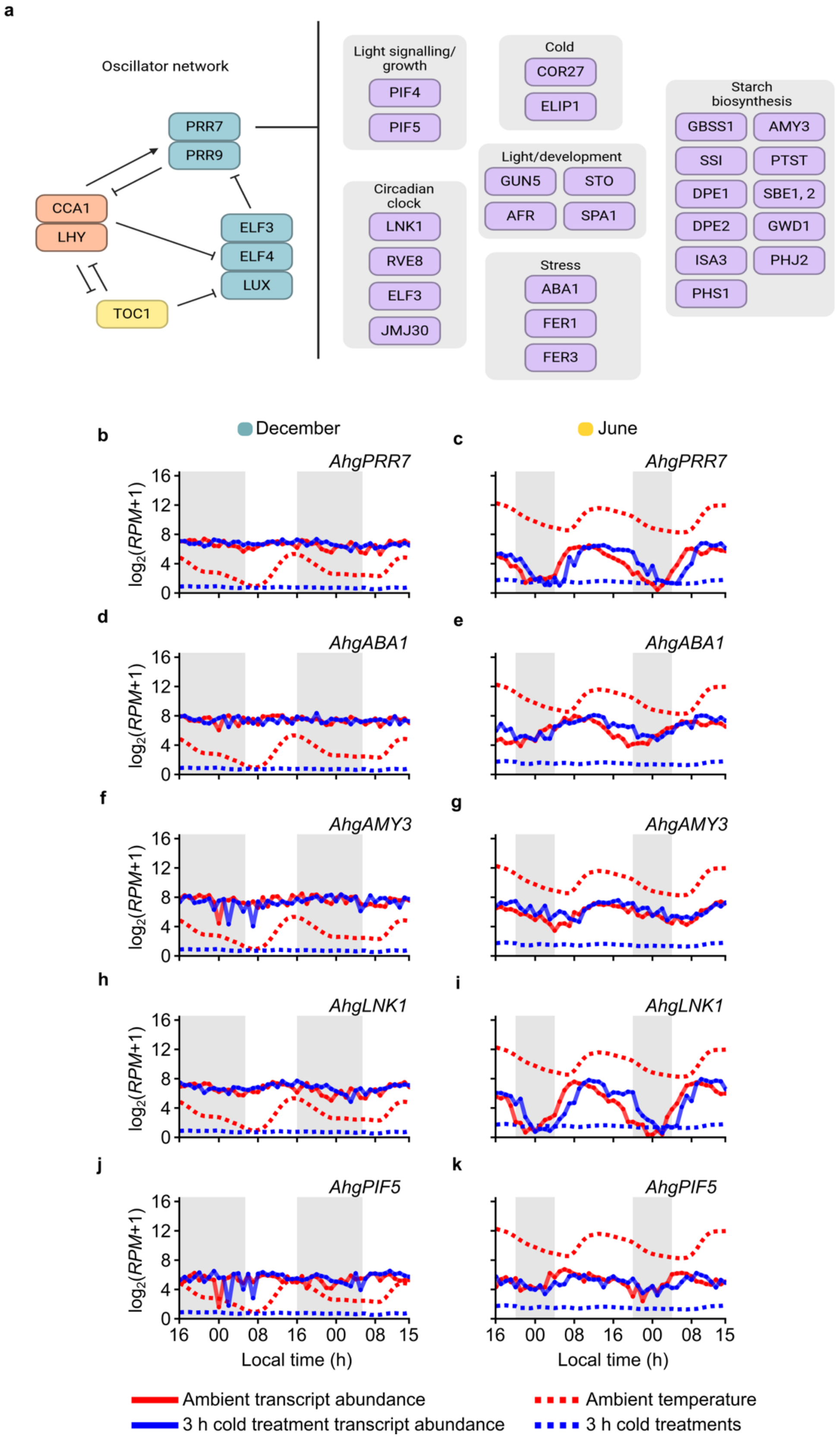


**Figure S8.** Temporal relationship between a clock component and its regulatory targets under naturally fluctuating conditions. This example considers the case of AhgPRR7. (a)

Simplified diagram of parts of the circadian oscillator network from *A. thaliana* (left) and some identified regulatory targets of PRR7 in *A. thaliana* (right, purple). (b-j) For two sampling seasons, time-series of (b, c) *AhgPRR7* transcripts and its regulatory targets (d, e) AhgABA1, (f, g) AhgAMY3, (h, i) AhgLNK1, a circadian oscillator associated component, and (j, k) AhgPIF5. See Discussion for explanations.

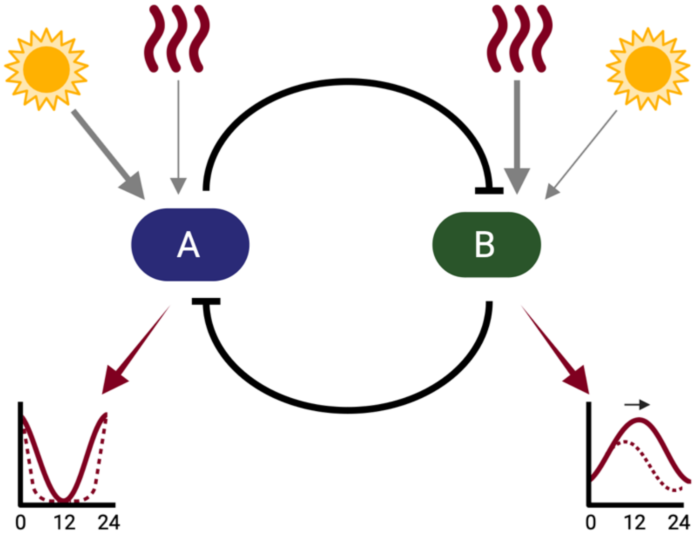


**Figure S9.** Concept of the gating of circadian clock outputs in the presence of competing zeitgebers. The circadian clock integrates various environmental cues that it receives, using these either for entrainment, to gate environmental signalling, or both. During the morning, Component A receives light and temperature information. For this clock component, light information is the prevailing zeitgeber, while clock outputs have temperature responses that are gated by the clock. The entrainment of Component A affects the circadian phase of Component B, which is entrained predominantly by temperature rather than light conditions, in turn affecting Component A until this is again regulated by light. Both Component A and Component B can be involved in circadian gating of the environmental responses of other transcripts, where the roles of Component A and Component B may differ in producing distinct gating classes. The hypothetical gating class of an output gene from Component A follows its response of A to the temperature cue (producing no clock phase shift), whereas the gating class of an output gene gated by Component B may receive the same phase shift as Component B does, due to its entrainment to the temperature cue. Sun and wave icons indicate light and temperature cues, respectively. Grey arrow thickness indicates the effectiveness of each cue in regulating each of Component A and B. Time-series diagrams at bottom indicate distinct gating classes.

**Supplementary Material References**

1 Romanowski, A., Schlaen, R. G., Perez-Santangelo, S., Mancini, E. & Yanovsky, M. J. Global transcriptome analysis reveals circadian control of splicing events in Arabidopsis thaliana. *The Plant Journal* **103**, 889-902 (2020). https://doi.org:https://doi.org/10.1111/tpj.14776
